# How low temperatures affect long-tailed bat (*Chalinolobus tuberculatus*) and lesser short-tailed bat (*Mystacina tuberculata*) activity, central North Island, New Zealand

**DOI:** 10.1101/2025.08.13.670222

**Authors:** Elyse Hyde, Stephanie S. Godfrey, Kerry M. Borkin

**Affiliations:** Wildlife Management, Department of Zoology, University of Otago; Senior Lecturer, Department of Zoology, University of Otago; Fauna Science, Terrestrial Biodiversity, Department of Conservation, Taupō

**Keywords:** survey effectiveness, *Mystacina tuberculata rhyacobia*, winter, temperature threshold, central lesser short-tailed bat

## Abstract

Bat activity generally increases with temperature, so bat surveys are considered more effective in summer. Less is known about survey effectiveness at low temperatures and whether activity ceases under certain thresholds. Literature provides piecemeal information on winter activity of threatened New Zealand lesser short-tailed bats (*Mystacina tuberculata*) and long-tailed bats (*Chalinolobus tuberculatus*), no data exist on lower temperature thresholds below which activity is halted in some regions. We recorded bat echolocation calls over seven nights in winter 2021 to determine whether a lower temperature threshold exists where both species are inactive in the Pikiariki Ecological Area, Pureora Forest Park, central North Island, Aotearoa New Zealand. Long-tailed bat detection rates greatly reduced at <5 °C, ceasing at <2.6 °C; at edge sites, long-tailed bats had higher activity rates at higher temperatures, but this pattern did not hold in the forest interior. Lesser short-tailed bats were detected at all sampled temperatures, (1 °C–10 °C), 56% of detections occurred at the forest edge; no lower temperature threshold beyond which activity ceased was found. While winter surveys may detect bats, rates are likely lower than in warmer months. Lesser short-tailed bat surveys could be worthwhile in winter due to their broader activity range, depending on the survey’s objectives, but should be conducted over extended periods, and any conclusions about the presence or absence of bats made cautiously. Further research is needed to refine lower temperature thresholds and improve winter survey effectiveness.

## Introduction

Surveys for bats – checks to see whether bats are present in a defined area – typically involve using acoustic recording devices (detectors) that detect bats’ echolocation calls. Understanding when or when not to survey for bats is essential. If surveys occur at a time of year or temperature when bats are inactive, results may indicate an absence of bats, even if they are present in the area. Bats’ activity drivers include weather conditions, metabolic demands, intraspecific competition, predation pressures and physiological adaptations (Borkin et al. 2023, Christie 2006, McNab and O’Donnell 2018, O’Donnell 2000, Speakman et al. 2003). Despite temperature regularly emerging as a common influence on species activity (in Aotearoa New Zealand, O’Donnell 2000; as well as in Europe, Laurence et al. 2025; Australia, Turbill 2008; and North America, Wolbert et al. 2014), activity patterns of bats during winter are considered perhaps one of the least documented of all their annual behaviours (Laurence et al. 2025).

Mainland Aotearoa New Zealand (henceforth Aotearoa) has two extant threatened bat species (pekapeka in te reo Māori, the indigenous language of Aotearoa: long-tailed bats (*Chalinolobus tuberculatus*) and lesser short-tailed bats (*Mystacina tuberculata*)); another species may persist on offshore islands (O’Donnell et al. 2023). Lesser short-tailed bats are most often detected using cluttered areas of the forest interior, whereas long-tailed bats are often most detected using forest edges and open spaces (O’Donnell et al. 2006). Both use torpor to reduce energetic costs. Torpor can vary from one-day to multi-day bouts, and whether lesser short-tailed bats arouse from torpor can be related to the temperature at sunset (Czenze et al. 2017). Both take shelter (roost) during the day, and parts of the night; often in large cavities within hollowed trees and tree trunks (O’Donnell and Borkin 2021, Parsons and Toth 2021). When not emitting ultrasound, such as when they are in torpor, they cannot be detected using acoustic recording devices.

Temperature strongly influences the detection rates of both species, with both more likely to be detected at higher temperatures (Borkin et al. 2023, Christie and Simpson 2006). However, being active in winter may be less energetically expensive for lesser short-tailed bats than for long-tailed bats because of their larger size, which may reduce heat loss, and ability to access a wider range of foods (O’Donnell and Borkin 2021, Parsons and Toth 2021, Thomas 2004). Lesser short-tailed bats can fold their wings to act as front limbs enabling foraging in leaf litter on the forest floor for invertebrates, and access to nectar or fruit (Kirsch et al. 1998, O’Donnell et al. 2021). Meanwhile, the long-tailed bat is largely restricted to aerial invertebrate prey, whose availability is markedly reduced in lower temperatures (O’Donnell 2000, 2001).

Differences between habitat use, morphology and ecology may result in variations in detection rates of Aotearoa’s bat species. Lesser short-tailed bats are known to be active at temperatures as low as −1.4 °C in Fiordland, southern Aotearoa (Sedgeley 2001). Long-tailed bats often do not emerge from their roosts when temperatures are <5 °C (Griffiths 2007), but they have been observed flying at lower temperatures (−1.5 °C; O’Donnell 2000).

Whilst guidance exists for other forest and habitat types in the central North Island of Aotearoa (Borkin et al. 2023), and more broadly throughout Aotearoa (e.g., see O’Donnell 2000 and others), there is no specific guidance in the literature about when acoustic surveys for the threatened bats of Aotearoa should take place in the relatively cool podocarp-hardwood forests of the central North Island, and at what temperatures bats are active there throughout winter, and, therefore, are available to be detected. No data are available on lower thresholds for either species ceasing activity completely. Despite this, bat ecologists are often asked whether surveys during winter are worthwhile because the detectors usually used for these surveys are more likely to be available at this time of year compared to in summer – the generally accepted time of year when surveys (inventories for the presence of bats) and monitoring (which looks for patterns in activity by repeated periods of recording with additional rigour) take place (KMB, obs.).

With this in mind, we wanted to understand whether there was a relationship between activity patterns – as measured by the number of call sequences – and temperature for each bat species, and particularly to understand whether there were temperature thresholds below which bats were not recorded. In this pilot study, we examine the influence of temperature on bat activity during winter by recording bat activity over seven nights in the 2021 winter season. We hypothesised that lower temperatures will be associated with reduced bat activity. We also explore the relationship between activity and detection rates in the forest interior and edge, and bat activity patterns throughout the night. We investigated the activity patterns of bats during winter at the Pikiariki Ecological Area in Pureora Forest Park, central North Island, Waikato, New Zealand with the intent of providing advice around surveying for bats at lower temperatures.

## Materials and Methods

### Study Area

This study was conducted within Pikiariki Ecological Area in Pureora Forest Park, central North Island, Waikato, New Zealand (38°31’01.2’S, 175°34’48.0’E), 549 m above sea level and > 78,000 ha (Dennis and Gartrell, 2015). Both lesser short-tailed bats (‘central’ subspecies, *M. t. rhyacobia*) and long-tailed bats roost nearby in this forest. We conducted our study in austral winter, from 2/08/2021–9/08/2021, to maximise the chance that low, potentially torpor-inducing, temperatures were included in the survey period.

### Sampling Design

To detect bat echolocation calls, we used six pairs of detectors (Frequency Compression Automated Bat Monitoring Units, AR4, Department of Conservation, Wellington).

Within each pair, one detector was placed at the forest edge, and the other within the forest c.60 m from its pair at the edge. This placement was to help ensure bats echolocating at one site would be unlikely to be detected at the other, so could be considered relatively independent (60 m is greater than the 30–50 m detection distances estimated for AR4 detectors (Smith et al. 2020) with forest in-between likely also increasing attenuation rates). Each pair was then placed 200 m apart to establish relative independence between pairs. Unfortunately, one detector (from site 7, group 4) malfunctioned during the study period, so we discarded data collected from that pair (group 4) to maintain a balanced study design. Several pairs of detectors were within 500 m of communal lesser short-tailed bat roosts used over the past summer (1/12/2020–31/2/2021, K Borkin pers. Obs.). Detectors were set to record from half an hour before sunset to half an hour after sunrise on nights beginning 2^nd^ to 8^th^ August 2021. This encompasses the times each species emerges from their roosts in the central North Island (K. Borkin, pers. Obs., Feng et al. 2025).

The call files recorded on each detector were processed using BatSearch v3.23 software (Department of Conservation, Wellington, New Zealand) and recordings were manually classified as either long-tailed bat or lesser short-tailed bat calls. We defined a call file as containing a call sequence when there were one or more bat calls in a spectrogram.

Weather data (temperature on the hour, °C) were downloaded from the nearest CLiFLo weather station to the study site (CLiFLo: Pureora Forest CWS, station 37016).

Data were organised to represent seven overnight periods (5 p.m.–7 a.m.). We allocated each call sequence to an hourly interval and identified the lowest hourly temperatures at which calls of each species were detected. The time of call intervals was converted into hours after sunset (1–14 hours after sunset, representing 5 pm–7 am) for further analysis.

### Data Analysis

To evaluate the effect of temperature and time of night on bat activity, while allowing for the estimation of non-linear relationships between these variables, we used a generalise additive mixed model (GAMM). Bat activity (number of bat call sequences per hour) was included as the response variable, modelled as a negative binomial distribution (family = ‘nb’) due to the skewed distribution of call data. Hours after sunset and temperature were included as smoothed variables, while location was included as a categorical variable. Night and block (detector pairs) were included as random effects to account for the non-independence of observations recorded on the same nights and within detector pairs. In addition, to investigate the relationship location had with the predictor variables and bat activity, a tensor product interaction (te) was added between location and temperature, and location and hours after sunset. Q-Q plots and residual plots were examined to ensure that data met the model assumptions using the functions appraise() and gam.check() in the package mgcv (Wood 2004). All analyses and data visualisation were carried out using R (version 4.4.1, R Core Team 2024).

## Results

### Temperature and bat activity

Over the study period, temperatures ranged between 1.3 °C and 10 °C, and it snowed on the final night of data collection. We recorded 2262 long-tailed bat call sequences and 424 lesser short-tailed bat call sequences. Long-tailed bats were detected at temperatures of 2.6 °C–10 °C, with 90% of detections recorded above 6 °C, and only one detection at the lowest temperature (Figure 1A). We recorded temperatures lower than 2.6 °C on four nights (the second, third, fourth, and seventh nights). We recorded long-tailed bat call sequences on each night of our study, but not after 9:54 p.m. on the last night when it snowed. Lesser short-tailed bats were recorded across the entire range of temperatures experienced during our study period (1.3°C-10°C). Lesser short-tailed bat call sequences were not recorded on the final night of the study, when it snowed.

**Figure 1.**
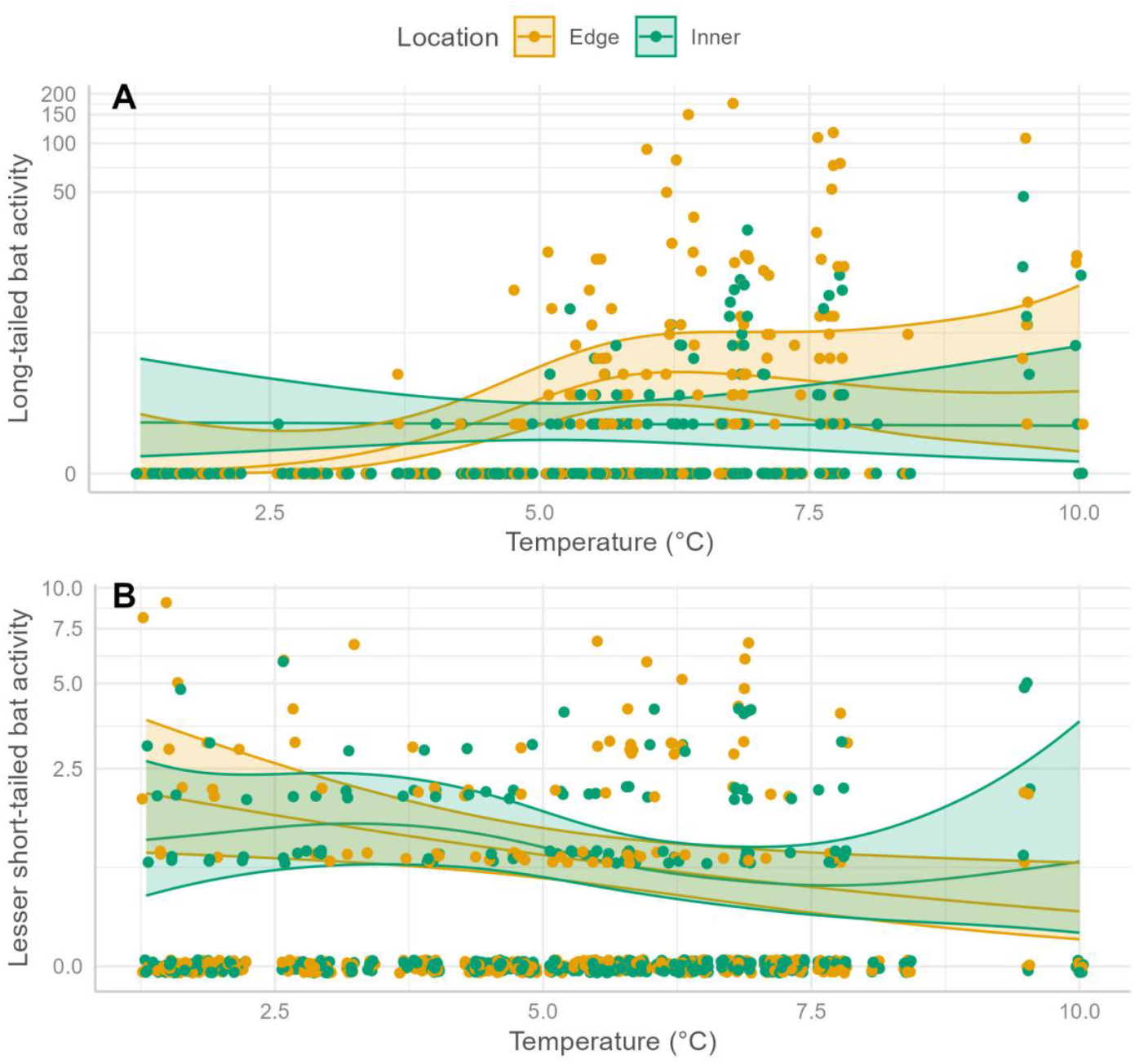
The relationship between number of call sequences per hour (bat activity) and temperature (°C) at edge and inner habitat locations of (A) long-tailed bats, and (B) lesser short-tailed bats in the Pikiariki Ecological Area, central North Island, New Zealand. Lines represent smooth estimates from the generalised additive mixed model, with shaded area representing 95% confidence intervals of the smoothed estimate. The y-axis is presented on a log scale.

Long-tailed bat call sequence detection rates varied with temperature for edge locations, with higher rates at higher temperatures (*X*^2^ = 10.80, edf = 3.16, *P* = 0.036); but not at inner locations where rates were independent of temperature (*X*^2^ = 0.004, edf = 1, *P* = 0.952, Figure 1A). Lesser short-tailed bat call sequence detection rates also varied with temperature for edge locations, where they declined linearly with temperature (*X*^2^ =5.66, edf = 1, *P* = 0.018), but rates did not vary with temperature at inner locations (*X*^2^ =5.61, edf = 2.85, *P* = 0.185, Figure 1B).

### Hours After Sunset and Bat Activity

For both species, detection rates varied through the night. The hour after sunset significantly influenced long-tailed bat activity at both edge (*X*^2^ = 118.28, edf = 7.36 *P* = <0.001) and inner locations (*X*^2^ = 79.18, edf = 5.30, *P* < 0.001). At both locations, call sequences per hour were highest at sunset, and remained high until 5 hours after sunset when rates began to decline, ceasing at 14 hours after sunset (Figure 2A). Lesser short-tailed bat call sequence detection rates also varied with time (hours) after sunset at both edge (*X*^2^ = 21.38, edf = 4.37, *P* = 0.001) and inner locations (*X*^2^ = 32.60, edf = 6.49, *P* = 0.001). Both edge and inner locations showed a similar relationship between hours after sunset and detection rate (Figure 2B). Lesser short-tailed bats were recorded over a longer period of the night than long-tailed bats (i.e., between 6 p.m.-6 a.m.). The first lesser short-tailed bat detection occurred two hours after sunset and peaked at 4–5 hours after sunset. Activity generally declined through the night from this peak; albeit with two smaller peaks in activity at eight hours after sunset (12 a.m.) and then after 13 hours (5 a.m.).

**Figure 2.**
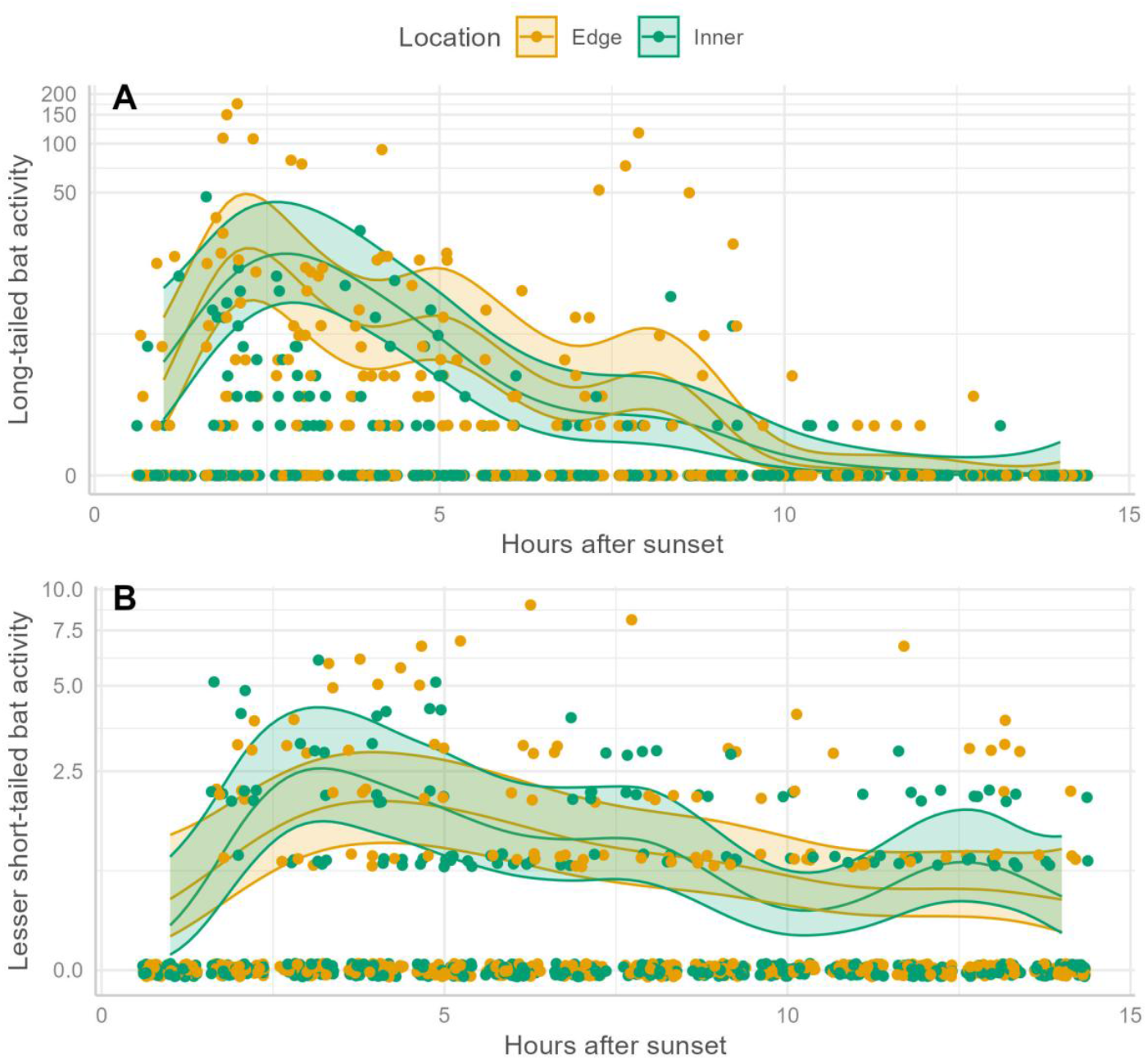
The relationship between number of call sequences (bat activity) and the hours after sunset at edge and inner locations of (A) long-tailed bats, and (B) lesser short-tailed bats in the Pikiariki Ecological Area, central North Island, New Zealand. Lines represent smooth estimates from the generalised additive mixed model, with shaded area representing 95% confidence intervals of the smoothed estimate. The y-axis is presented on a log scale.

## Discussion

This study provides the first interrogation of winter bat activity in the relatively cool podocarp-hardwood forests of the central North Island, and the first for the central subspecies of the lesser short-tailed bat. As expected from other studies (e.g., Borkin et al. 2023, O’Donnell 2000), we recorded more long-tailed bat call sequences at higher temperatures, with only 10% of calls occurring below 6 °C. This is unsurprising as long-tailed bats do not always emerge from roosts when sunset temperatures are less than 5 °C (Griffiths 2007). Whilst the number of nights we recorded call sequences for was small (*n* = 7), we detected no long-tailed bats at temperatures <2.6 °C. Further south, in Fiordland’s Eglinton Valley, long-tailed bats still fly at lower temperatures (−1.5 °C, O’Donnell 2000). Whether 2.6 °C reflects a threshold for long-tailed bat activity in the central North Island requires further data collection at a wider range of temperatures. The results reinforce the findings of others (O’Donnell 2000, Borkin et al. 2023) that surveys at low temperatures (in our case, < 6 °C) are less likely to detect long-tailed bats. Supporting this, Borkin et al. (2023) found that long-tailed bats were most likely to be detected between 8 °C and 17 °C in a nearby exotic forest, with fewer detections at colder temperatures.

We recorded lesser short-tailed bats at all temperatures they experienced during our study period. Our results align with other studies that suggest that temperature has less influence over lesser short-tailed bat activity rates than it does for long-tailed bats (O’Donnell 2000, O’Donnell et al. 2006, Borkin et al. 2023). This may be due to lesser short-tailed bats’ lower reliance on aerial invertebrates, broader diet, and slightly higher body mass (O’Donnell and Borkin 2021, O’Donnell et al. 2021, O’Donnell et al. 2006, Thomas 2004). Despite this, Christie and Simpson (2006) found that in southern New Zealand, the likelihood of lesser short-tailed bat detections in winter increased with every 2.2 °C rise in overnight mean temperature. As other researchers have suggested, we predict that while lesser short-tailed bats have been observed flying at −2 °C in southern New Zealand (Sedgeley 2001), their lower temperature thresholds may be higher in warmer northern climates.

Long-tailed bats might be more constrained by temperature than lesser short-tailed bats due to a combination of their diet and morphology. Whilst minimum temperature likely determines whether bats emerge from torpor and are available to be detected (Czenze et al. 2017), invertebrate activity is thought to strongly influence foraging duration (O’Donnell 2000) and therefore the number of detections. This may be particularly so when the species is heavily reliant on flying invertebrates for most of their food.

At our study site, long-tailed bats eat mainly aerial invertebrates (Diptera comprised 40% of items in faeces, Lepidoptera 24%, and Coleoptera 18%, Gurau 2014), as do lesser short-tailed bats (Czenze et al. 2018). As temperatures decrease, arthropods experience physiological restraints such as reduced metabolic rates, locomotion and feeding, inhibiting activity in low temperatures (Stejskal et al. 2019). Invertebrates that walk are more likely to be active at lower temperatures than those that fly and so are still available as prey at lower temperatures (Stejskal et al. 2019). This suggests that bats that feed on non-flying invertebrates (such as lesser short-tailed bats), will have food available for longer periods each night and for a greater proportion of nights, because their prey are active over a wider range of temperatures. When their diet also includes fruit, nectar and pollen as do lesser short-tailed bats (Parsons and Toth 2021), they may be even less restricted by temperature. Lesser short-tailed bats have slightly higher body mass than long-tailed bats which may also allow them to more easily maintain their body temperature (Fenton et al. 2003). This could explain why we found lesser short-tailed bats were active both at lower temperatures, and over a longer period of the night, than long-tailed bats in our study.

Patterns of activity differed between long-tailed bats and lesser short-tailed bats. We observed a peak in activity an hour after sunset (1–2 h after sunset) for long-tailed bats, after which activity was greatly reduced. Long-tailed bats were rarely recorded in the last three hours of the night. In contrast, detection rates varied less throughout the night for lesser short-tailed bats. Lesser short-tailed bat activity was first recorded and peaked an hour later than long-tailed bats, and they were detected over more of the night. It is important to acknowledge that temperature and hours after sunset are interrelated (i.e. as the night progresses, usually temperature decreases), thus it is impossible to truly separate these two variables when interpreting these patterns.

Activity patterns throughout the evening could partially be driven by temperature, especially at the start and end of the evening, when temperatures are at their greatest extreme. Regardless, we suggest that this later peak and longer period of activity for lesser short-tailed bats reflects their later emergence and lesser reliance on flying invertebrates for food. Surveys for bats need to take these differences into account by ensuring periods when bats are most likely to be active are included.

On the final night of our study, snow fell in the Pureora region. We detected no lesser short-tailed bat call sequences on that night, but recorded long-tailed bats earlier that evening (last calls were recorded at 9:54 p.m. during that seventh night). Snow likely fell from 2 a.m. onwards, as earlier temperatures remained between 4.4 °C and 6.5 °C. The complete absence of lesser short-tailed bat detections that night suggests that another factor in addition to temperature influences their activity. This absence was particularly notable given detectors were within 500 m of known communal roosts used over the summer period. This highlights the risk of very short surveys not detecting bats when present. Further research is needed to investigate which other weather variables affect lesser short-tailed bat detection rates, and the effect of microclimatic differences on both species.

We recorded long-tailed bats most frequently at the forest edge, typical of other studies (O’Donnell et al. 2006). The inner forest detections exhibited similar activity peaks at lesser intensities as the outer forest, although inner activity ceased after six hours. Although typically associated with the forest interior (O’Donnell et al. 2006, Parsons and Toth 2021), we recorded lesser short-tailed bats using both forest edges and interiors at similar detection rates. Whilst this may be because in the forest interior calls attenuate over shorter distances compared to those emitted along a forest edge due to the presence of clutter (Scrimgeour et al. 2013), it highlights to surveyors that detectors placed at the forest edge may still detect lesser short-tailed bats.

Surveys for lesser short-tailed bats may be worthwhile undertaking in winter, if they cannot be completed during a warmer time of year, because they appear to be detected at a wider range of temperatures than are long-tailed bats. Surveying for long-tailed bats at the same time will increase survey efficiency. However, surveys in winter should take place over a long period to ensure that nights with higher temperatures are included, and should be interpreted with caution, taking into account the risk of not detecting bats when present. We recommend continuing research aiming to understand lower temperature thresholds for lesser short-tailed bats and long-tailed bats in this region, and to understand by what quantum surveys during winter are less likely to detect bats. Surveys that are relied upon for identifying whether trees have bats present in them, or which decide on which/whether management action takes place, should not take place at such low temperatures because of the risk that bats are not detected when present.

## Acknowledgements

We thank mana whenua for their support of this research, Maniapoto Area office of the Department of Conservation for their support during field work, and the Threatened Species Research Workstream of the Department of Conservation for equipment.

Specific funding was not received for this research.

